# Peripheral Inflammation Limits Serotonin Neuron Signaling Capacity via Serotonergic IL-1R1 to Reduce Neuronal Excitability and Enhance Serotonin Clearance

**DOI:** 10.1101/2025.10.13.682078

**Authors:** P. A. Gajewski, H. Iwamoto, A. N. Tillman, Z. Filliben, A. E. Walsh, N. L. Baganz, M. J. Robson, M. Zapata, N. Quan, R. D. Blakely

## Abstract

Neurobehavioral disorders, ranging from depression to schizophrenia, have been found to display immune system alterations. The high incidence of comorbidity of these disorders, particularly depression, with chronic inflammatory conditions suggests shared mechanisms contributing to the manifestation of these conditions. We have previously shown that peripheral modulation of the innate immune system in mice rapidly triggers enhanced serotonin (5-HT) clearance *in vivo* associated with increased anxiety- and despair-like behaviors that can be suppressed by serotonergic elimination of p38α MAPK. Forebrain-projecting 5-HT synthesizing neurons of the dorsal raphe nucleus (DRN^5-HT^) play a key role in regulating behaviors related to mood and anxiety and whose perturbations are observed in multiple affective disorders. Here we identify molecular and circuit-level mechanisms that can translate peripheral innate immune system activation into changes in 5-HT signaling capacity. Using whole cell patch clamp recordings from acute midbrain slices, we demonstrate that the proinflammatory cytokine, IL-1β, acts cell autonomously through its receptor, IL-1R1, via the p38α MAPK signaling pathway to rapidly inhibit firing of DRN^5-HT^ neurons. In the dorsal hippocampus, we found that as with acute, peripheral lipopolysaccharide (LPS) administration, local injections of IL-1β rapidly enhance 5-HT clearance as assessed by *in vivo* chronoamperometry. Like IL-1β, TNFα also acts via a serotonergic p38α MAPK dependent pathway to reduce excitability of DRN^5-HT^ neurons. Immunocytochemical studies reveal that, IL-1R1, is nonuniformly expressed by DRN^5-HT^ neurons and is required for LPS-induced inhibition of these cells as detected by cFos activation, with sex-dependent patterns evident. Moreover, we detected both DRN^5-HT^ IL-1R1-dependent and -independent LPS-mediated changes in cFos changes in forbrain projection areas. Our findings support a growing appreciation that serotoninergic neurons contribute to changes in CNS physiology and behavior following peripheral immune activation. More specifically, our studies attest to a functional role of serotonergic IL-1R1 in mediating IL-1β following peripheral innate immune activation, effects likely to arise both from changes in diminished 5-HT neuron excitability and elevated 5-HT clearance.

## Introduction

Infection by a pathogenic microbe results in an immediate response by the body’s innate immune system. One of the main effects of activating the innate immune system is a coordinated and complex engagement of various immune cells to remove the pathogen and any infected host cells. The signals that activate and recruit these cells are largely the province of peptides known as proinflammatory cytokines, and as such, elevated levels of these molecules in the bloodstream of an individual is a typical sign of infection (1). Once released, inflammatory cytokines circulate widely, including through the brain due to its extensive vascularization, where an echo of peripheral immune activation can be transmitted into the parenchyma by activation of endothelial cytokine receptors (2). Cytokines can also be produced directly in the brain by cells known as microglia, although levels here are relatively small compared to the periphery and typically only increase in response to brain trauma or direct central nervous system (CNS) infection (3, 4). One particular cytokine, interleukin-1 beta (IL-1β) is thought to initiate an amplifying cascade of a number of other proinflammatory cytokines and neuroinflammation-associated molecules (5). High, prolonged levels of IL-1β produce sickness behavior, manifested as fatigue, loss of appetite, irritability, and social withdrawal, among other highly conserved behavioral features that are presumed to reflect an organism’s attempt to conserve energy, allow for bodily processes to restore homeostasis, and limit the spread of infection to others. Specific brain regions including the hypothalamus are key regulators of sickness behavior and known to be modulated by inflammatory cytokines (6). However, most neuronal populations lack high level expression of the requisite receptors to respond directly. Recent evidence indicates serotonin (5-HT) synthesizing neurons of the DRN (DRN^5-HT^) are a unique population of neurons within the midbrain that are under the influence of various cytokines, most notably IL-1β (7–9). 5-HT, via the wide-ranging projections of both the DRN^5-HT^ and the more medial raphe nuclei (MRN^5-HT^) regulate a multitude of behaviors including those influenced by chronic innate immune activation. Importantly, elevated levels of inflammatory cytokines have been found in association with multiple mood disorders (10, 11). Similarly, chronic inflammatory conditions, whether produced in a disease state with immune activation, such as rheumatoid arthritis, or intentionally through medications as a byproduct, such as interferon-α, are associated with depression (12, 13). Thus, even in the absence of an active infection, peripheral elevations of inflammatory cytokines are still able to influence behavior, many of which are impacted by serotonergic signaling.

One mechanistic explanation often used to support the neuroinflammatory hypothesis of major depressive disorder proposes a role for elevated inflammatory cytokines in altering 5-HT metabolism and signaling in the CNS (14). Neuropsychiatric disorders such as depression, anxiety, obsessive compulsive disorder, autism spectrum disorder, and schizophrenia have long been associated with dysfunction in 5-HT signaling, reinforced by the use of medications that specifically inhibit the presynaptic 5-HT transporter (SERT), a plasma membrane protein responsible for shaping the magnitude and duration of 5-HT signaling following release. SERT activity and density *in vitro* and *in vivo* is under intricate regulation from multiple pathways, including several contributing to innate immune responses. In addition to the impact that inflammatory cytokines have on 5-HT metabolic pathways (15), activation of cytokine receptors, such as interleukin-1 receptor type 1 (IL-1R1), has been shown to increase SERT intrinsic activity via activation of p38α MAPK, predicted to decrease the extracellular levels of 5-HT available for signaling (9, 16, 17). At the somatic level, there is also evidence that IL-1β has an inhibitory effect on the excitability of 5-HT neurons, though whether effects derive from actions directly on serotonergic neurons themselves or their afferents, or both is debated (18, 19). Regardless, these findings support the contention that inflammation decreases 5-HT signaling, driving expression of sickness-related behaviors, that when sustained are underlying facets of the behavioral changes found in neurobehavioral disorders. Interestingly, it also appears that treatment with selective serotonin reuptake inhibitors (SSRIs) have an effect on the levels of peripheral cytokines, likely due to the presence of SERT on circulating platelets and the role that 5-HT has in the peripheral immune system in addition to its role as a central neurotransmitter (20–22). Microglia also express 5-HT receptors, allowing for crosstalk in the CNS between 5-HT signaling and locally produced inflammatory cytokines (23).

Here we investigate the role of serotonergic IL-1R1 in modulating 5-HT neuronal activity by using acute systemic inflammatory stimuli as well as direct cytokine application to *ex vivo* brain slices. We find a nonuniform distribution of serotonergic IL-1R1 among DRN^5-HT^ neurons and that these neurons are inhibited in response to inflammatory cytokines IL-1β and TNFα. Additionally, a peripheral inflammatory stimuli causes serotonergic IL-1R1-dependent changes in neuronal activity not only within the DRN^5-HT^ neuronal population as well as in downstream projection sites. Moreover, acute peripheral inflammation alters the transcriptional profiles of mouse DRN^5-HT^ neurons in a sex-dependent manner, revealing unique responses of these cells in males and females.

## Methods

### Animals

All housing, breeding, and procedures/experiments were performed according to the NIH Guide for the Care and Use of Experimental Animals and approved by the Florida Atlantic University Animal Care and Use Committee (IACUC). Mice were housed with 12-hour light cycles with food/water *ad libitum*, and tissue was harvested during the light period. A 5-HT neuron-specific deletion of IL-1R1, ePet1:Cre;IL-1R1^loxP/loxP^ was used to investigate the necessity of serotonergic IL-1R1 in mediating a response to peripheral inflammation. A 5-HT neuron-specific restoration of IL-1R1, ePet1:Cre;IL-1R1^r/r^ was used for the visualization of serotonergic neurons expressing IL-1R1 via a transcriptional reporter (tdTomato) inserted into the endogenous *il1r1* gene as well as an HA-tag fused to the IL-1R1 protein (24). Whole-cell patch-clamp recordings were performed on adult ePet1:EYFP mice, a generous gift from Dr. Evan Deneris (25). Both male and female mice, age 8-12 weeks, were used.

### Lipopolysaccharide treatment

To induce innate immune system activation, we administered lipopolysaccharide (LPS, intraperitoneal injection of 0.2 mg/kg, 026:B6, Sigma, #L8274) or 0.9% sterile saline at 10 ml/g body weight. Three hours after treatment, mice were sacrificed by transcardial perfusion (methods noted below) and used for either immunohistochemistry or spatial transcriptomics.

### Electrophysiological recordings

Whole cell patch clamp recordings were performed on acute midbrain slices containing the DRN, with serotonergic neurons detected via direct fluorescence imaging of EYFP expressed by ePet1. Brains were sliced coronally into 280 *μ*m thick sections with a vibratome (VT1000 S, Leica Biosystems) in ice-cold carbonated (95% O_2_ and 5% CO_2_ mixture) sucrose-substituted artificial cerebrospinal fluid (87mM NaCl, 75mM sucrose, 2.5mM KCl, 1.25mM NaH_2_PO_4_, 25mM glucose, 7mM MgCl_2_, 0.5mM CaCl_2_, 25mM NaHCO_3_). After recovery for 30 min in the same solution at 28°C, the slices were equilibrated in artificial cerebrospinal fluid (ACSF) (124mM NaCl, 2.5mM KCl, 1mM NaH_2_PO4, 2.5mM CaCl_2_, 1.3mM MgSO_4_, 26mM NaHCO_3_, 20mM glucose, 310 mOsm/kg) for a hour at room temperature. Slices were submerged in a chamber (RC-27, Warner instruments) mounted on a fixed-stage upright microscope (Axioskop 2 FS, Zeiss) equipped with infrared differential interference contrast (IR-DIC) optics. IR-DIC images through 40x water-immersion objective were visualized with a cooled CCD camera (ORCA R2, Hamamatsu) controlled by HCImage software (Hamamatsu). ACSF was perfused with a peristaltic pump through in-line heater (28°C, TC-324B, Warner Instruments) at a rate of 1 mL/min and complete exchange of the recording chamber volume occurred in approximately 3 min. Recording electrodes with 5-7 MΩ were prepared from a capillary glass (8250, A-M systems) on a micropipette puller (P-97, Sutter Instrument). Internal pipette solution was composed of 135 mM K gluconate, 10 mM KCl, 10 mM HEPES, 1 mM EGTA, 1 mM MgATP, 0.2 mM Na_2_GTP, 1 mM Na_2_ phosphocreatine, 2 mM MgCl_2_, pH 7.3 adjusted with KOH, 295 mOsm/kg. Whole-cell current clamp recordings were made using MultiClamp 700B amplifier (Molecular Devices). Currents were low pass filtered at 2 kHz and digitized at 5 kHz with Axon Digidata 1550B (Molecular Devices) and stored using pClamp 10.7 software (Molecular Devices). Series resistance was not compensated. Access resistance was limited to less than 20 MΩ and recordings with larger than 20 MΩ were discarded. 0.5-1.0 Hz of action potentials were generated by adjusting current clamp value. The changes in firing rates with IL-1β perfusion were calculated using Clampfit software. For reversal potential measurements, 1 sec of 20 mV step-voltages were applied from +130 to −30 mV under voltage clamp in presence of 0.5 µM tetrodotoxin. Membrane potential was not corrected for liquid junction potential, which is estimated to be −15 mV (Clampex junction potential calculator). Recombinant mouse IL-1β and TNFα were obtained from R&D systems (#201-LB-005/CF and #10291-TA-020). p38α MAPK-selective inhibitor MW150 was a generous gift from Dr. D. Martin Watterson, Northwestern University Feinberg School of Medicine (26).

### Synaptosomal uptake

[^3^H] 5-HT uptake was measured in synaptosomes prepared from the midbrain as previously described (27). Assays were conducted in 1-mL Krebs–Ringer’s HEPES assay buffer (containing 130mM NaCl, 1.3mM KCl, 2.2mM CaCl_2_, 1.2mM MgSO_4_, 1.2mM KH_2_PO_4_, 1.8 g/L glucose, 10mM HEPES, pH 7.4, 100 μM pargyline and 100 μM ascorbic acid). After assessment of protein levels (Bradford assay, Bio-Rad #5000201), 20–30 μg synaptosomes per sample (in a total volume of 200 μl) were pre-incubated at 37°C in a shaking water bath for 5–10 min. Modifiers were then added for 10 min, and samples were incubated with 20 nM [^3^H] 5-HT 5 min at 37°C. Uptake was terminated by adding 1 ml ice-cold Krebs–Ringer’s HEPES buffer, followedby filtration through GF/B Whatman filters that had beensoaked in 0.3% polyethylenimine for 1 hr before experiments. Trapped radioactivity was eluted in scintillation fluid (Ecoscint H, National Diagnositics) overnight and quantified by scintillation spectrometry. Specific counts were obtained after subtraction of counts obtained from parallel samples assayed in the presence of 10 μM paroxetine.

### *In vivo* chronoamperometry

Recordings of exogenously applied 5-HT clearance were obtained from the CA3 subregion of the dorsal hippocampus (−1.94mm A/P, +2.0mm M/L, −2.0mm D/V from bregma) as previously described (28). Under the conditions used, basal 5-HT clearance is primarily mediated by SERT (29). Once reproducible 5-HT signals were obtained, IL-1β (2 ng) was administered locally 30 min prior to administration of 1mM 5-HT to the same location. Recording sites were verified postmortem by histological analysis of hippocampus sections.

### Fluorescence *in situ* hybridization (FISH)

Mice were anesthetized using a fatal dose of Euthasol™ and perfused transcardially with ice cold 1x PBS followed by ice cold 4% paraformaldehyde (PFA, pH 7.4) in phosphate buffered saline (PBS), pH 7.4. Brains were removed, postfixed in 4% PFA for 24 hrs and then equilibrated in a cryoprotective solution of 30% sucrose in PBS at 4°C for 24-48 hrs. FISH was conducted via RNAScope multiplex version 2 via per approaches recommended by Advanced Cell Diagnostics (ACD, document number: 47-007-CKL). Slices were incubated in a mixture of channel 1 (*il1r1*, 488161) channel 2 (*Tph2*, 318691-C2) and channel 3 (Tnfrsf1a, 426541-C3 or Tnfrsf1b, 435941-C3) probes pipetted directly onto each section until fully submerged. The HRP-C1, C2, and C3 signals were developed by incubating in Opal dyes (Akoya Biosciences, diluted 1:1500 in the RNAscope TSA buffer). After the final wash, excess liquid was removed from the slides and RNAscope supplied DAPI (4′,6-diamidino-2-phenylindole) was added. DAPI was removed and Prolong Gold Antifade Mountant (Invitrogen, #P36961) was added and coverslipped. Slides were imaged on a Nikon A1 confocal microscope at 60x magnification. Images were examined for Tph2-positive cells and presence of puncta to indicate the various immune receptor transcripts.

### Immunohistochemistry

Mice were anesthetized using a fatal dose of Euthasol™ and perfused transcardially with ice cold 1x PBS followed by ice cold 4% paraformaldehyde (pH 7.4) in PBS for our cFos labeling. Brains were removed, postfixed in 4% PFA for 24 hours and then equilibrated in a cryoprotective solution of 30% sucrose in PBS at 4□C for 24-48 hrs. Alternatively, for our HA and SERT co-labeling, mice were perfused with room temperature 1x PBS followed by room temperature 9% glyoxal and 8% acetic acid (GAA, pH 4.0) and postfixed in GAA for 24 hours before being placed in 30% sucrose until slicing. Brains were then sectioned on the freezing microtome into 35 mm thick coronal sections and stored in 1x PBS with 0.02% sodium azide until use. Sections were washed in 1x PBS with 0.1% Triton X-100, blocked with 3% normal donkey serum in 1x PBS with 0.2% Triton X-100. Afterwards, slices were incubated with primary antibodies: rabbit anti-RFP (Abcam #ab185921, 1:1000), goat anti-Tph2 (Immunostar #22941, 1:1000), rabbit anti-cFos (Abcam #ab302667, 1:2000), guinea pig anti-SERT (Frontiers #MSFR103300, 1:500), and rabbit anti-HA (Cell Signaling #3724, 1:1000). Primary antibody incubations were completed overnight at 4°C (except for anti-HA, which was incubated at room temp overnight). After washing to remove excess primary antibody, secondary antibodies were added (donkey anti-rabbit Cy3 used at 1:500, Jackson Immunoresearch #711-165-152; donkey anti-goat Alexa Fluor 488 used at 1:500, Jackson Immunoresearch #705-545-003; donkey anti-rabbit HRP used at 1:500, Jackson Immunoresearch #711-035-152). cFos-only sections (lateral habenula and central amygdala) underwent diaminobenzidine (DAB) labeling (Vector laboratories, SK-4100). Sections were mounted on Superfrost plus slides (Fisher) and dried. Mounted slides were dehydrated in an ethanol gradient and cleared using Citrisolv before being cover-slipped using DPX mountant (distyrene, plasticizer, and xylene; Sigma #06522). Representative images were captured using a Nikon A1R confocal microscope. Multiple-channel images were overlayed when necessary and cells were counted by a blinded observer manually by first identifying 5-HT-synthesizing neurons (Tph2-positive) and then counting the number of cFos-positive nuclei among them. DAB labeled cFos was automatically counted using the NIS Nikon Elements Analysis software with the following parameters: 15-120 (low-high) intensity, size (10-20 px), circulatiry (0.3-1.0).

### GeoMx DSP Assay

Whole brains were mounted on a freezing microtome (Leica) to slice 8 µm sections that were promptly mounted on Superfrost plus slides (Fisher Scientific, #12-550-15) and allowed to dry at room temp for 24 hrs before being stored at −80°C until use (no more than 2 weeks later). All subsequent steps were performed using RNase-free conditioned and DEPC treated water. Slides were removed from the freezer, allowed to warm to room temp before being incubated at 60°C for 1 hr. Slides were then washed in 50%, 70%, and 100% ethanol for 5 min each, air dried for 10 min and then antigen retrieval performed using target retrieval reagent (Invitrogen, #00-4956058) for 15 min at 95°C. Slides were then washed in 1X PBS and incubated in 0.1 mg/ml proteinase K (Invitrogen, #AM2546) for 15 min at 37°C and washed again at room temp. Slides were incubated with whole mouse transcriptome atlas hybridization probes diluted in Buffer R (provided in the GeoMx RNA Slide Prep FFPE PCLN kit and NGS RNA WTA mM, #121300313 and #121401103) in a hybridization oven at 37°C for 18 hrs. Following probe incubation, slides were washed with stringent washes (equal parts formamide and 4XSSC buffer (Invitrogen, #AM9763)) 2X at 37°C for 25 min. Then slides were washed 2X in 2X SSC buffer. Slides were incubated in buffer W (provided in the GeoMx RNA Slide Prep Kit) for 30 min and incubated in morphology marker for Tph2 (goat anti-Tph2, 1:100, Abcam, #ab1210313) in buffer W for 2 hrs at room temp. Slides were washed in 2X SSC buffer and then incubated with a secondary antibody (donkey anti-goat 647, 1:200, Jackson Immunoresearch, #705-605-003) and the nuclei marker Syto13 (GeoMx nuclear stain morphology kit, #999181) diluted in buffer W. Syto13 immunofluorescence was used for the autofocus of GeoMx imaging. Immunofluorescence for Tph2 was used to identify DRN^5-HT^ neurons. Each region was located approximately −4.72mm A/P from bregma. The DRN was divided into a dorsal and ventral portion (see Fig 1). The dorsal portion contained the dorsal and dorsolateral wing subregions, whereas the ventral portion contained the ventral and interfascicular subregions. Illumina NextSeq 1000 was used for sequencing with a read length of 27 for both reads with reverse sequence orientation in the readout group plate information. Plates were dried and rehydrated in 10 mL of nuclease-free water, mixed, and incubated at room temp for 10 min. PCR was performed on samples as described in the Nanostring GeoMx DSP Readout User Manual using 2 ml PCR master mix, 4 mL primer from the correct wells, and 4 mL resuspended DSP aspirate. Libraries were pooled and sequenced. Fastq files were converted to digital count conversion (DCC) files using GeoMx NGS Pipeline x2.3.3.10. Sequencing reads were trimmed to 27 bp to reflect GeoMx probe length. In the GeoMx experiment, 24 segments were collected in total from 12 mice and all segments passed Quality Control (QC) analysis. Of the 20,175 gene targets contained in the GeoMx Mouse Whole Transcriptome Atlas, 8581 were included in the downstream analyses following QC analysis. To account for systemic variation between areas of illumination (AOIs), we normalized each count by the 75^th^ percentile (Q3) of expression for each AOI. To ensure normalization and QC procedures were accurate, we compared each AOI’s Q3 value to its limit of quantification. Differential expression of gene targets in the GeoMx experiment was determined via t-test model which held two of our three variables (subregion, genotype, or treatment) constant while assessing differences in the third variable. Gene names identified from the volcano plot analysis were then input into Enrichr software (Developed by the Ma’ayan Lab, Icahn School of Medicine at Mount Sinai, Center of Bioinformatics) for gene ontology (GO) biological processes pathway analysis (30–32).

**Figure 1.**
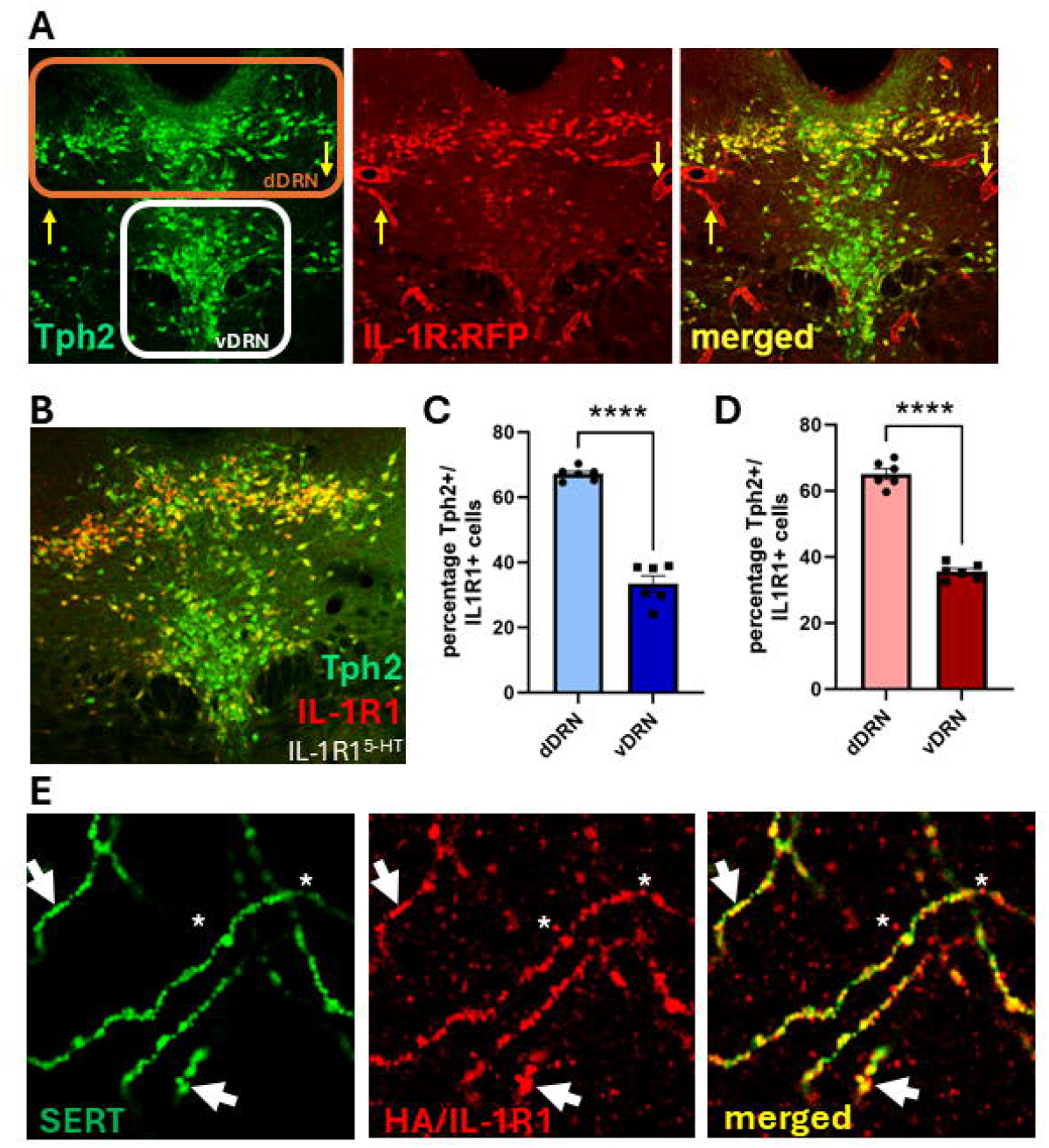
Serotonergic IL-1R1 expression in DRN^5-HT^ neurons predominates in dorsal and dorsolateral DRN. A. Representative images from a male IL-1R1 global restore mouse (24) within the DRN showing 5-HT synthesizing (Tph2) neurons in green and IL-1R1 expression tracked by tdTomato in red (IL1R:RFP). The orange box highlights the dorsolateral and dorsal subregions of the DRN (dDRN) whereas the white box highlights the ventral and interfascicular subregions of the DRN (vDRN) where relatively fewer IL-1R1-positive serotonergic neurons are found. Yellow arrows indicate endothelial expression of IL-1R1. B. Representative image from the DRN of a male ePet1:Cre;IL-1R1^r/r^ mouse where IL-1R1 expression is restricted to serotonergic neurons. Tph2 antibody labels 5-HT neurons (green) and a transcriptional reporter (RFP) defines the expression of IL-1R1 proteins (red). C. Percentage of IL-1R1-positive cells within male DRN^5-HT^ neurons comparing the dorsal and ventral subregions. The dDRN shows a significantly higher percentage of serotonin-positive, IL-1R1-positive neurons compared to the vDRN. D. Percentage of 5-HT-positive, IL-1R1-positive cells within the female DRN comparing the dorsal and ventral subregions. The dDRN shows a significantly higher percentage of serotonin-positive, IL-1R1-positive neurons compared to the vDRN. E. Representative image from male dorsal hippocampus of serotonergic fiber labeled with SERT (green) and the HA-tagged IL-1R1 (red) from an ePet1:Cre;IL-1R1^r/r^ mouse. White arrows indicate colocalized SERT and IL-1R1 along a serotonergic axon. * indicates instances of IL-1R1 labeling along a serotonergic axon without SERT. Unpaired t-test ****p < 0.0001 dDRN versus vDRN.

### Statistical analysis

Statistical analyses, comparing male versus female, saline versus drug-induced changes in IHC, were performed with GraphPad Prism 10. T-tests, two-way analyses of variance (ANOVA) with subsequent planned comparisons (Bonferroni) are noted in the legends. Electrophysiology data were analyzed with Clampfit 10.7 (Molecular Devices), OriginPro 2017 (OriginLab), and Prism 6 (GraphPad).

## Results

### Neuroanatomical characterization of serotonergic IL-1R1

The DRN^5-HT^ neurons are found in distinct subregions of the mouse midbrain and display unique projection targets. By examining the distribution of *il1r1* expression through the DRN subregions, we can determine a subset of downstream brain regions likely impacted by the activation of serotonergic IL-1R1. To localize IL-1R1^+^ DRN^5-HT^ neurons, we utilized an *Il1r1* reporter line affording a transcriptional tdTomato (RFP) reporter and an HA epitope tag translational reporter that can be used to localize sites of IL-1R1 protein expression (24). Both reporters are under the control of the endogenous *il1r1* promoter and mobilizable under the control of Cre recombinase. A globally restored *il1r1* mouse expresses tdTomato within any cell type that endogenously expresses *il1r1*. Figure 1A shows expression of RFP throughout various cell types within the DRN including Tph2-positive cells and endothelial cells lining local blood vessels. Previous findings have revealed GABAergic neurons within the DRN also to express *il1r1*, and indeed we detected limited cell body RFP within the DRN that was not Tph2-positive in this mouse line, suggesting that these may be GABAergic DRN neurons (7). We then limited *il1r1* expression to DRN^5-HT^ neurons by crossing the IL-1R1^r/r^ reporter line with ePet1-Cre to better characterize the distribution of serotonergic IL-1R1 neurons within the DRN. We found a distinct difference between the dorsal and ventral subregions of the DRN in numbers of RFP (IL-1R1 expressing) labeled cells that were double labeled with anti-Tph2. Both the males and females show higher levels of expression in the dorsal and dorsolateral wing subregions (collectively termed the dorsal DRN; 65% colocaliation) compared to the ventral and interfascicular subregions (collectively termed the ventral DRN; 35% colocalization) (Figure 1B-D), although no differences existed in the percentage of DRN^5-HT^ neurons expressing IL-1R1 between males and females. This difference in DRN^5-HT^ IL-1R1 distribution likely impacts downstream signaling, as each of these subregions of DRN^5-HT^ neurons are known to project to unique brain regions. In one key projection region, the dorsal hippocampus, we sought evidence of IL-1R1 on serotonergic fibers, combining SERT labeling with visualization of IL-1R1 via an in frame fusion of an HA epitope, with serotonergic specifiy of expression achieved our ePet1:Cre;IL-1R1^r/r^ mouse line. We observed extensive colocalization between SERT and IL-1R1 along serotonergic axon fibers with overlap evident at varicosities identified by the beaded structure of axons (Figure 1E). In some fields, we also observed a number of IL-1R1 positive fibers lacking SERT expression. This adjacent IL-1R1 may reflect pools of IL-1R1 being translocated down axons to distal sites or indicate that SERT is more specifically localized near sites of 5-HT release.

### IL-1β suppresses excitability of DRN^5-HT^ neurons in acute brain slices

Brain-wide mapping of neuronal IL-1R1 expression has recently been achieved by Nemeth et al., revealing prominent populations of neuronal receptor expressions in the hippocampus, sensory cortex, and raphe nuclei (8). Multiple studies have implicated IL-1β in reducing activity of DRN^5-HT^ neurons, although neither the specificity of IL-1R1 involvement nor intracellular pathways activated have been interrogated (18, 19). When we perfused acute midbrain slices with IL-1β (10 ng/mL) while monitoring activity of DRN^5-HT^ neurons with whole cell patch clamp, we observed a reliable decrease in the frequency of firing of these cells accompanied by a hyperpolarization (Figure 2A, B). To confirm that the decrease in firing rate of the DRN^5-HT^ neurons was cell autonomous and not via expression of IL-1R1 on other nearby cell types, we repeated these recordings using slices from ePet1:Cre;IL-1R1^r/r^ mice where *il1r1* expression is driven only in DRN^5-HT^ neurons. Whole cell patch clamp recordings yielded responses similar to those obtained using a wildtype background (Figure 2C), supporting the conclusion that expression of IL-1R1 on other cell types in the slice is not involved in IL-1β action on DRN^5-HT^ neurons. Consistent with this conclusion, recordings obtained in the absence of inhibitors for Glu/GABA signaling failed to demonstrate a loss of IL-1β effects (data not shown). Supporting the specificity of IL-1β effects, pretreatment of slices with the IL-1R1 antagonist (IL-1ra) resulted in a loss of IL-1β inhibition of DRN^5-HT^ neuron firing (Figure 2D). Prior studies have reported that IL-1β increases the intrinsic activity of SERT in a p38α MAPK-dependent manner *in vitro* (9, 16). Moreover, loss of serotonergic p38α MAPK limits the effect of peripheral lipopolysaccharide (LPS) on SERT-mediated 5-HT uptake in midbrain synaptosomes as well as despair and anxiety-like behavior (17). We thus determined whether the inhibitory actions of IL-1β on DRN^5-HT^ neurons were p38α MAPK dependent. Consistent with our prior synaptosomal studies, the actions of IL-1β on DRN^5-HT^ neurons were significantly attenuated by the highly-specific p38α MAPK inhibitor MW150 (17, 26, 33) (Figure 2E, F). The voltage dependence of IL-1R1 mediated current of DRN^5-HT^ neurons demonstrates a nonlinear profile with a reversal potential at −83 mV (Figure 2G) consistent with an increase in K^+^ channel conductance as mediating the reduction in firing frequency. We next used a genetic approach to investigate endogenous cytokine actions by examining midbrain synaptosomes derived from mice with a selective IL-1R1 elimination in serotonergic neurons (ePet1:Cre;IL-1R1^loxP/loxP^) and treated with the bacterial mimetic, LPS. These effects were found to be consistent with an essential role of IL-1R1 in mediating serotonergic responses to inflammation as the LPS-mediated stimulation of 5-HT uptake was no longer present when mice lacked serotonergic expression of *il1r1* (Figure 2H). Finally, we utilized *in vivo* chronoamperometry (34) in the presence of local IL-1β, examining the cytokine’s effects on a key forebrain projection, the hippocampus, where we previously found systemic LPS to induce a rapid, time-dependent stimulation of 5-HT clearance (16). As predicted, local injection of 2ng IL-1β within the CA3 subregion produced a significantly faster 5-HT clearance rate compared to vehicle control (Figure 2I). Last, we explored whether the inhibitory action on DRN^5-HT^ neuron excitability can be observed with slice application of TNFα, another proinflammatory cytokine we previously documented to stimulate SERT activity in cultured cells *in vitro* (9). Consistent with our prior *in vitro* findings, bath perfusion of TNFα (10 ng/mL) reduced the firing rate of DRN^5-HT^ neurons in midbrain slices, effects that are blocked by MW150 (Figure 3A, B), as similarly seen with IL-1β. We then utilized FISH to examine the expression of TNFα receptors by DRN^5-HT^ neurons as there are multiple TNFα receptors with unique intracellular signaling pathways. We confirmed the presence of TNFαR2 (*tnfrsf1b*) on DRN^5-HT^ neurons whereas TNFαR1 (*tnfrsf1a*) did not localize to 5-HT synthesizing neurons, as identified by *tph2* in situ (Figure 3C, D). Importantly, the TNFαR2 appears on the same DRN^5-HT^ neurons also expressing *il1r1*, which can be seen with our triple label in situ (Figure 3C).

**Figure 2.**
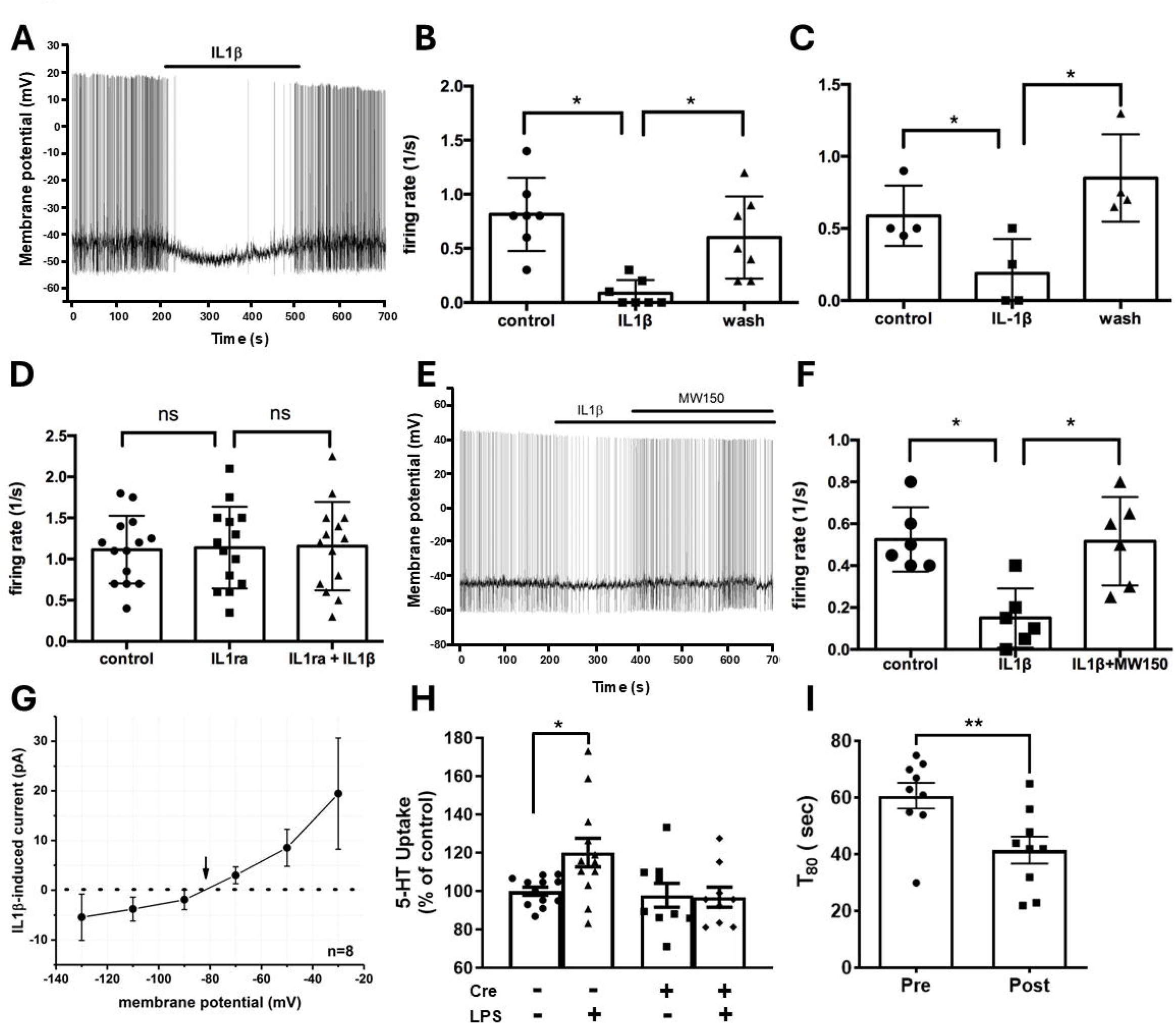
IL-1*β* reduces DRN^5-HT^ excitability via a p38*α* MAPK dependent mechanism and enhances 5-HT uptake *ex vivo* and 5-HT clearance *in vivo*. A. Current clamp recordings from DRN^5-HT^ neurons demonstrate inhibition of excitability produced by IL-1β. Recordings were obtained using elicited action potentials before and after bath perfusion of IL-1β (10ng/mL). Neural activity was inhibited and accompanied by hyperpolarization of the membrane potential. B. Quantitation of impact of IL-1β on DRN^5-HT^ neurons. Averaged firing rates from individual cell recordings. First period of the recording (0-200s) was used as control. IL-1β effect was calculated between 200-500s, prior to washout. C. Quantitation of effect of IL-1β on DRN slices from ePet1:Cre;IL1R1^r/r^ mice reveals sufficiency of serotonergic IL-1R1 expression to reduce excitability of DRN^5-HT^ neurons. Averaged firing rates were calculated. First period of the recording (0-200s) was used as control. IL-1β effects were calculated between 200-500s, prior to washout. D. IL-1R1 antagonist (IL-1ra) inhibits IL-1β-induced reductions in raphe neuron excitability. IL-1ra (10ng/ml) perfused alone had no effect on the average firing rate but eliminated the ability of IL-1β (10ng/ml) to reduce DRN^5-HT^ neuron excitability. E. IL-1b effects on DRN^5-HT^ neuron excitability are mediated via p38α MAPK signaling. Current clamp recordings were obtained using elicited action potentials before and after bath perfusion of IL-1β (10ng/ml). Neural activity was inhibited and accompanied by hyperpolarization of membrane potential that could be rescued upon bath perfusion of the p38α MAPK inhibitor, MW150 (1uM). F. Quantification of excitability impact of IL-1β and subsequent MW150 treatment. First period of the recording (0-200s) was used as control. IL-1β effect was calculated between 200-500s, prior to MW150 treatment. G. IV relationship of IL-1β induced current of DRN^5-HT^ neurons. Current-voltage (IV) curve derived from voltage steps before and after IL-1β treatment demonstrated a non-linear effect on membrane potential. Reversal potential was −83mV (indicated by arrow), consistent with activation of potassium channels. H. Elimination of serotonergic IL-1R1 expression (ePet1:Cre;IL-1R1^loxP/loxP^) resulted in the elimination of the LPS-induced increase in midbrain synaptosomal [^3^H]5-HT 5-HT uptake. Two-way ANOVA, followed by post hoc Tukey’s multiple comparison test * p < 0.05, demonstrated 5-HT uptake stimulation in synaptosomes prepared from ePet1:Cre negative mice administered LPS versus vehicle that could be abolished in synaptosomes prepared from ePet1:Cre positive mice. I. Enhanced rate of 5-HT clearance following in vivo IL-1β injection. Amperometric recordings were obtained in the CA3 subregion of the dorsal hippocampus of wildtype C57 male mice. Local IL-1β (2ng) was administered, and the rate of clearance was measured as the time it takes 80 percent of 1mM 5-HT to be cleared. “Pre” and “post” signify clearance times of 5-HT before and after local injections of IL-1β respectively. **p < 0.01, Pre vs Post.

**Figure 3.**
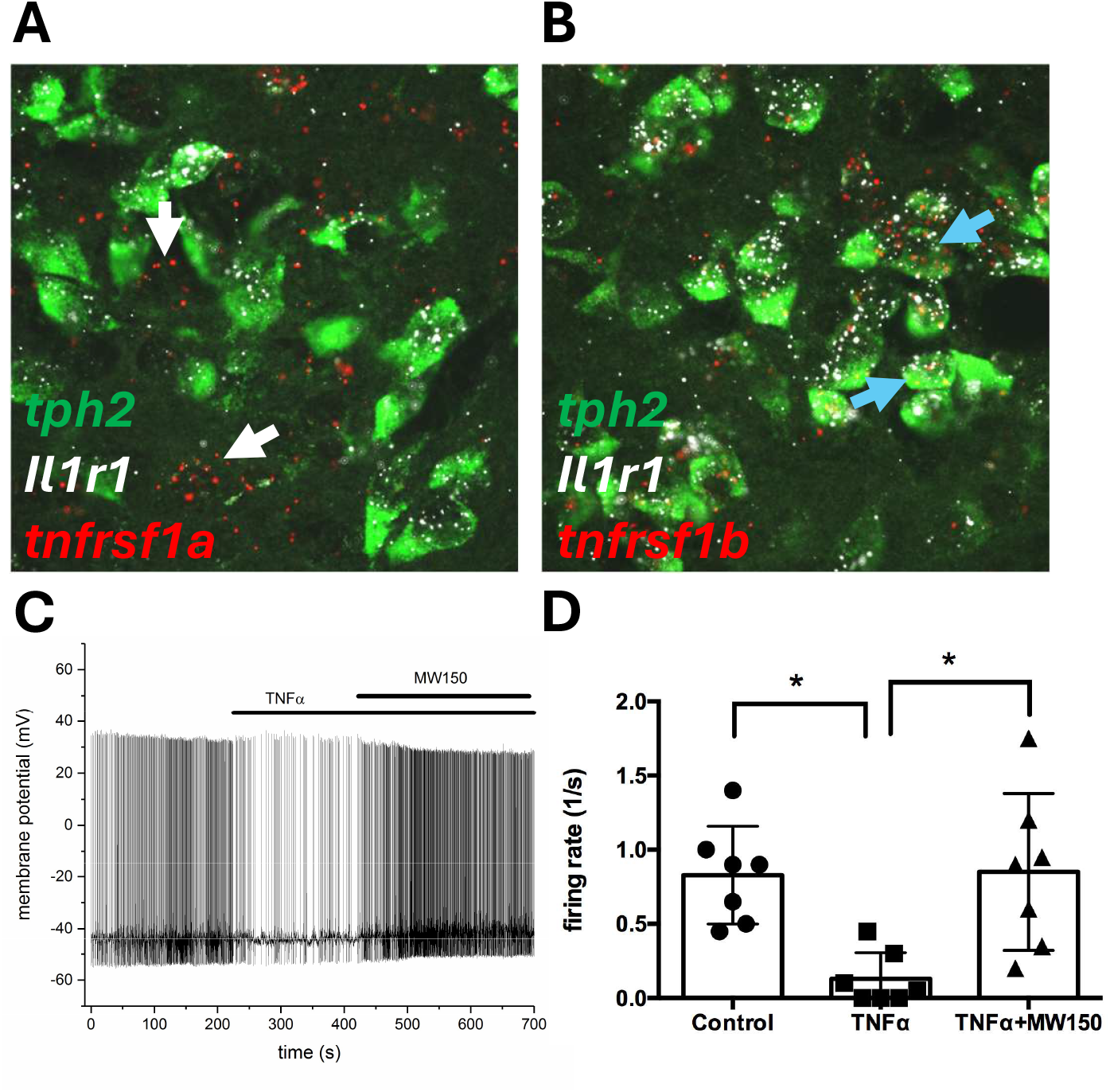
TNF*α* receptor expression on DRN^5-HT^ neurons and electrophysiological effects of TNF*α* on DRN^5-HT^ neuron excitability. A. Current clamp recordings were obtained using elicited action potentials before and after bath perfusion of TNFα (10ng/ml). Neuronal activity was inhibited and accompanied by hyperpolarization of resting membrane potential. B. Quantification of excitability impact of TNFα and subsequent MW150 treatment. First period of the recording (0-200s) was used as control. TNFα effect was calculated between 200-500s, prior to MW150 treatment. C. Representative image of *in situ* hybridization (RNAscope) within the dorsal DRN. Tissue was probed for Tph2 (green), Il1r1 (white), and Tnfrsf1a (red). No Tph2-positive cells were found to express Tnfrsf1a. White arrows indicate Tnfrsf1a-positive cells that are non-serotonergic. D. Representative image of FISH within the dorsal DRN. Tissue was probed for Tph2 (green), Il1r1 (white), and Tnfrsf1b (red). Multiple Tph2-positive cells were found to co-express Il1r1 and Tnfrsf1b. Blue arrows indicate serotonergic neurons expressing Tnfrsf1b.

### Acute, low dose LPS reduces DRN^5-HT^ activity in a region, sex- and IL-1R1-dependent maner

Above, we established a cell autonomous IL-1R1 dependent suppression of DRN^5-HT^ excitability by IL-1β. Next, we sought to determine whether IL-1R1 also supports a change in neuronal activation of these cells in the context of a peripheral inflammatory stimulus, 0.2 µg/mL LPS, monitoring subsequent expression of the immediate early gene, cFos, in our ePet1:Cre;IL-1R1^loxP/loxP^ mice. The dose of LPS was chosen based on our prior work demonstrating a lack of sustained locomotor suppression typical of sickness syndrome and thus being relevant in our examination of mild systemic inflammation and not overt sickness behavior (16). We found that both male and female ePet1:Cre;IL-1R1^loxP/loxP^ (Cre+) mice, which are engineered to eliminate serotonergic IL-1R1, showed significantly fewer DRN^5-HT^/cFos^+^ co-expressing cells as compared to their Cre-littermates (Figure 4A), supporting a role of serotonergic IL-1R1 in mediating the effects after systemic LPS administration. We also observed a difference in median raphe nucleus 5-HT neurons (MRN^5-HT^) with a large portion of these cells being IL-1R1-positive (data not shown). Female Cre-mice show a similar LPS-mediated decrease in cFos-positive MRN^5-HT+^ neurons compared to DRN^5-HT^ cells. Interestingly, male MRN^5-HT^ cells did not display a significant decrease in cFos activation following LPS administration (Figure 4A). The serotonergic, IL-1R1-dependent LPS effect also appears to be subregion specific within the DRN of females, whereas males exhibit cells suppressed throughout the DRN (Figure 4B, C). Next, we examined forebrain projection regions that are known targets of DRN^5-HT^ neurons. Among the regions assessed, we found the lateral habenula to be uniquely impacted by serotonergic IL-1R1 signaling following peripheral LPS in females but not males (Figure 4D). LPS also induced neuronal activation in forebrain projection regions as shown by elevations in cFos^+^ cells in a sex- and serotonergic IL-1R1-independent manner in the central amygdala, a region known to show elevated neuronal activity in response to peripheral inflammation (35, 36) (Figure 4E).

**Figure 4.**
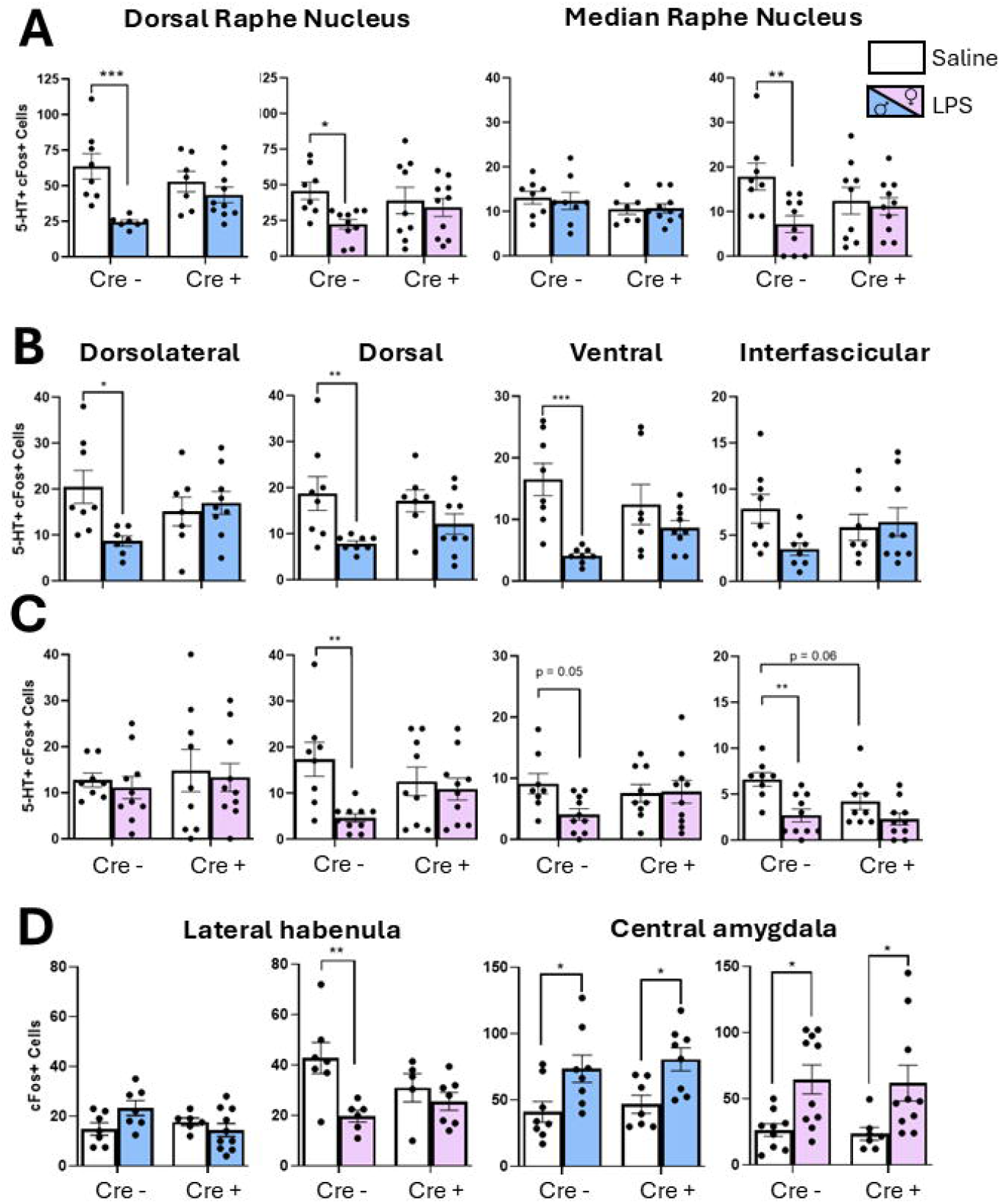
Acute peripheral administration of LPS impacts cFos labeling of DRN^5-HT^ neurons and DRN projection targets in an IL-1R1 and sex-dependent manner for. A. Quantification of 5-HT immunoreactive, cFos-positive cells within both the dorsal (left) and median (right) raphe nuclei in males and females of ePet1:Cre; IL-1R1^loxP/loxP^ mice. Cre negative (Cre-) mice maintain expression of serotonergic IL-1R1, whereas midbrain raphe 5-HT neurons of Cre positive (Cre+) mice lack the capacity to express IL-1R1. in. LPS administration (0.2mg/kg, i.p. 3hrs prior to sacrifice) caused a significant reduction in cFos-positive DRN^5-HT^ neurons of Cre^-^ males and females, whereas an inability to express IL-1R1 impaired the LPS-induced reduction of cFos-positive DRN^5-HT^ neurons. For female MRN^5-HT^ neurons, LPS also decreased the number of serotonergic cFos-positive nuclei in an IL-1R1-dependent manner whereas no impact of cFos was detected in males. B. Quantification of cFos-positive cells within DRN^5-HT^ neurons of each subregion of the DRN in male ePet1:Cre; IL-1R1^loxP/loxP^ mice. Nearly all subregions demonstrated a significant, LPS-induced reduction of cFos-positive 5-HT immunoreactive neurons (except for the interfascicular subregion) that was impaired in mice lacking the capacity to express serotonergic IL-1R1. C. Quantification of 5-HT-immunoreactive cFos-positive cells within each subregion the DRN in female IL-1R1^loxP/loxP^;ePet1:Cre mice. Only the dorsal subregion demonstrated a significant LPS-induced reduction of cFos-positive 5-HT neurons that is impaired with the loss of serotonergic IL-1R1. D. Quantification of cFos-positive nuclei within the lateral habenula (left) or central amygdala (right) after treatment with acute LPS in male and female IL-1R1^loxP/loxP^;ePet1:Cre mice. In the lateral habenula, female Cre-mice showed a significant decrease in cFos positive nuclei after LPS that was absent in mice with an inability to express serotonergic IL-1R1 (Cre+), whereas failed to show significant differences in cFos labeling in neither a Cre− or Cre+ context. In the central amygdala, male and female mice both demonstrated a serotonergic IL-1R1-independent increase in cFos positive nuclei in response to acute LPS. Two-way ANOVA, followed by post hoc Tukey’s multiple comparison tests, *p < 0.05, **p < 0.01, ***p < 0.001, saline versus LPS.

### DRN^5-HT^ neurons demonstrate sex-dependent gene expression differences *in situ* following acute LPS

The immediate physiological response to inflammatory cytokines we noted in our slice electrophysiology experiments are not likely derived from transcription dependent effects due to the transient properties of the stimulus and response. However, the observation that immediate early gene, cFos, appears to be altered three hours after peripheral LPS treatment indicates that peripheral inflammation may impart transcriptional changes to 5-HT neurons, given sufficient time. Given that cFos was found to demonstrate sex-dependent cFos activation in DRN^5-HT^ neurons following acute LPS administration, we asked whether transcriptional responses of these cells might demonstrate changes in a sex dependent manner as an initial response to a peripheral inflammatory stimulus. Thus we pursued spatial transcriptomic profiling of DRN^5-HT^ neurons, collecting samples within the DRN dorsal subregion that highly express IL-1R1 as well as the lower IL-1R1 expressing ventral regions (Figure 5A). We observed a sex-dependence in transcriptional responses with males having a significantly higher number of differentially expressed genes compared to females in response to LPS (Figure 5B, C). Of transcripts reduced by LPS compared to saline, only 2% of these were enriched in both males and females (Figure 5B). In the case of transcripts elevated by LPS compared to saline, less than 1.5% of them were similar between males and females (Figure 5C). The impact of peripheral LPS in the dDRN of male mice can be seen in the volcano plot in Figure 5D. LPS treatment in the males led to an enrichment of genes associated with synaptic development, structure, and plasticity (labeled Shank3, Adam10, and Rab5b in the volcano plot) (37–39) whereas genes diminished after LPS (higher in saline) associated with maintenance and protein synthesis (labeled Spcs2 and Akap7 in the volcano plot) (40, 41) (Figure 5D). To determine pathways dependent on DRN^5-HT^ IL-1R1 activation, we combined the list of genes reduced and elevated in response to LPS in both sexes to define a set of differentially expressed genes (DEGs) that are enriched in specific biological and physiological pathways. We performed gene ontology pathway analysis using the Enrichr program (see Methods) and observed that males and females lacked overlapping pathways, which based on their lack of overlapping DEGs, was not surprising. Overall, LPS appeared to generate DEGs in males associated with pathways linked to regulation of gene expression whereas in females, DEGs were more associated with regulation of synaptic function (Figure 5E).

**Figure 5.**
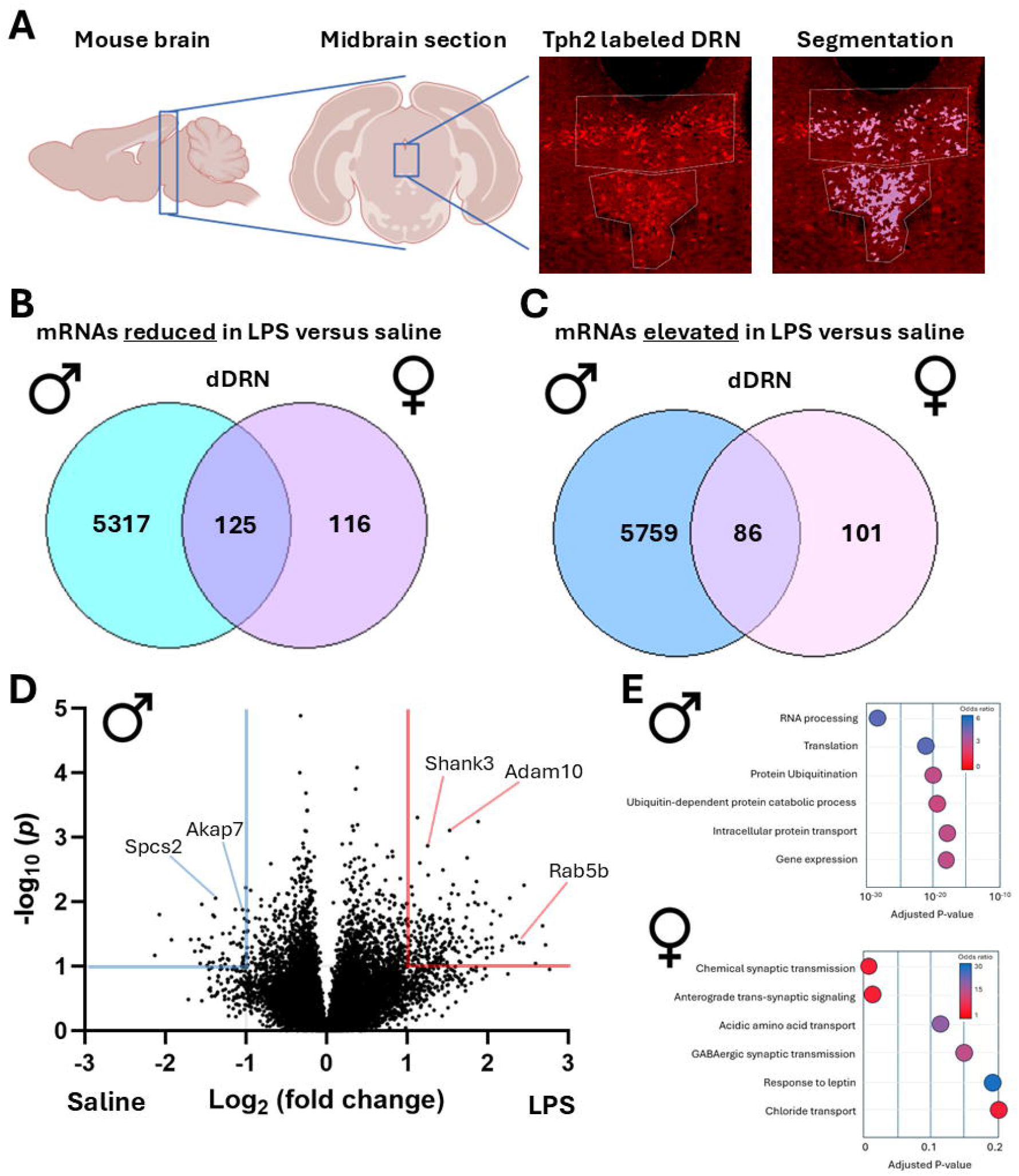
Peripheral LPS impacts transcriptional profile of DRN^5-HT^ neurons more in males compared to females. A. Diagram of mouse brain slice selection and identification of DRN^5-HT^ containing midbrain slices. Slice sections were labeled for Tph2 to identify 5-HT synthesizing neurons with subsections of the DRN taken for spatial trascriptomics analysis (ROIs identified by white boarders in image). Segmentation was performed based on Tph2 labeling to select only the serotonin neuron population for RNA sequencing. Diagrams were generated using Biorender. B. Venn diagram detailing an increased number of mRNA transcripts enriched in the male saline groups (reduced in the LPS treatment groups) as compared to females. C. Venn diagram detailing the total number of mRNA transcripts found enriched with LPS treatment (reduced in the saline treatment group). A minor portion of the reduced transcripts were found in both males and females. D. Volcano plot to display differential gene expression in the context of saline (negative log2) and LPS (positive log2) in DRN^5-HT^ neurons of the dDRN of male mice. D. Gene ontology term enrichment identified using Enrichr. Biological processes that include the involvement of the genes identified in B and C. Odds ratio refers to the strength of the association with that pathway.

## Discussion

Our studies document the regulation imposed by IL-1β on DRN^5-HT^ neuron excitability and the dependence of these effects on p38α MAPK signaling. These experiments highlight sex-specific transcriptome changes following acute systemic LPS, and changes in neuronal activation in projection areas as monitored by cFos expression. Generally speaking, IL-1β signaling to DRN^5-HT^ neurons is inhibitory, whether as a consequence of reducing neuronal excitability at the cell body level, demonstrated with *ex vivo* slice recordings, or by increasing 5-HT clearance, observed in serotonergic projection areas by *in vivo* chronoamperometry. Importantly, our peripheral LPS studies were performed at doses subthreshold for causing sickness behavior (16) in an effort to understand how the initial effects of inflammation target DRN^5-HT^ neurons in the absence of time-dependent alterations that arise with sickness models (42, 43).

As shown by genetic reporter and immunocytochemical studies, IL-1R1 appears to be nonuniformly expressed throughout subsets of DRN^5-HT^ neurons, with the highest level of expression found in the dorsal and dorsolateral wings, observed in both males and females. As subregions of the DRN are known to have distinct projection targets (44, 45), the varied levels of IL-1R1 expression throughout the DRN support a differential effect of IL-1β transmitted to forebrain targets. The dorsal subregion of DRN^5-HT^ neurons project to and influence regions related to stress reactivity (medial prefrontal cortex, central amygdala), reward and motivation (nucleus accumbens, caudate putamen, substantia nigra), pain modulation (rostral ventral medulla, central amygdala, medial prefrontal cortex), and behavioral and physiological flexibility (prefrontal cortex and the limbic system) (46–48). There is significant overlap between the dorsal and dorsolateral subregions in their projection targets, as both target the nucleus accumbens, paraventricular nucleus of the hypothalamus, central amygdala, and rostral ventral medulla. The dorsolateral wings also target regions related to sensory relay and integration (multiple thalamic nuclei), sensorimotor and attentional control (thalamic nucleus, superior colliculus, and ventrolateral orbital cortex), as well as spatial and contextual memory processing (entorhinal cortex and septum) (44). The lateral habenula, a region we uncovered as being uniquely affected by serotonergic IL-1R1 activation in a sex-dependent manner, receives dorsolateral DRN^5-HT^ innervation and typically uses inputs to encode aversive behaviors (49, 50). Affective-related brain regions are a known target for DRN^5-HT^ neurons and dysregulation there can result in anhedonia or fatigue. Such dysregulation of the serotonergic system may lead sensory systems to misfire and contribute to brain fog or hypersensitivity, and can disrupt general homeostatic circuits to produce disrupted sleep, appetite, and temperature control (51–53). Importantly, these symptoms of 5-HT dysfunction are behavioral symptoms of infection and to some extent, certain neuropsychiatric disorders such as depression.

This is to our knowledge the first direct detection of IL-1R1 protein expressed on serotonergic axons, made possible by the HA tag introduced in the IL-1R1^r/r^ mouse line (24). Additionally, greatly improved labeling of HA-IL-1R1 was achieved using glyoxal as a fixative over paraformaldehyde, as noted for other difficult to detect antigens (54). Visualization of IL-1R1 on serotonergic axonal fibers identified not only colocalization between IL-1R1 and SERT, but also adjacent IL-1R1 expression along the axons. Prior studies have noted a rapid effect that activation of IL-1R1 on SERT intrinsic activity which is likely made possible by their physical proximity (9, 16). Additional studies are needed to assess quantitatively the co-localization of SERT with IL-1R1 and a potential physical association that could account for the ability of IL-1β to regulate SERT activity.

As noted above, our finding that IL-1β enhances 5-HT clearance while acting at the cell soma level to limit DRN^5-HT^ neuron firing points to a coordinated and rapid response induced by the cytokine to diminish 5-HT signaling with the earliest signs of peripheral inflammation. It is possible that chronic immune activation may alter how cytokines can regulate activation and signaling of DRN^5-HT^ neurons. Although the effect of IL-1β to enhance SERT activity has been well documented in previous studies (9, 16, 17), the studies reported here are the first to demonstrate an increased ability of IL-1β to enhance 5-HT clearance *in vivo*. It is important to recognize that the slices used in our electrophysiological experiments are deafferented from afferent glutamatergic and GABAergic inputs and as such, although we obtain similar results in studies with slices bathed in antagonists of their receptors (data not shown), we cannot exclude that *in vivo*, IL-1R1 may regulate DRN^5-HT^ neurons via modulation of presynaptic inputs known to regulate these cells, possibilities that will require additional studies. Based on IV characteristics of IL-1R1 dependent current and the insensitivity of IL-1β effects to inhibitors of G-protein regulated inward rectifying (GIRK) channels (data not shown), IL-1β inhibition of DRN^5-HT^ neurons likely arises from activation of a to be identified potassium conductance sensitive to p38α MAPK-dependent activation. To date, limited studies that have investigated the relationship between p38 MAPK activation and potassium conductances. One such study found that p38 MAPK activation increased potassium channel activation upstream of caspase cleavage in the context of neuronal apoptosis (55). Others have found that p38 MAPK activation induced transcriptional changes of potassium channels (56), although due to the time frame of our electrophysiological experiments, such changes are unlikely to be in play. Importantly, all of the current electrophysiological experiments took place in the DRN subregions with the highest IL-1R1 expression as we sought to maximize our ability to monitor serotonergic IL-1R1 effects on DRN^5-HT^ neuron firing. As neither sex dependence nor subregion specificity were evaluated in these experiments, our understanding of whether these factors play a role in the serotonergic response to IL-1β (or TNFα), should be pursued in future studies. However, as our current electrophysiological studies did not indicate differences by sex in IL-1β response, this appears to be in line with the sex-independent actions of peripheral LPS on IL-1R1-dependent cFos reductions throughout the DRN.

Prior studies utilized *in vivo* chronoamperometry to reveal that peripheral LPS rapidly and transiently increases 5-HT clearance in the dorsal hippocampus (16). However, as noted, peripheral LPS causes a rapid upregulation of a multitude of cytokines. In order to confirm the effect of IL-1β on 5-HT clearance, we added IL-1β directly into the dorsal hippocampus (CA3 subregion) and observed similar effects to those found following peripheral LPS administration, though of course with a slower time course (16). As our *in vivo* chronoamperometry data were obtained using wildtype males, we cannot discount the likelihood of a sex-dependent effect of IL-1β effects on 5-HT clearance, warranting further study, as does demonstration that the IL-1β effects are IL-1R1 dependent, though our visualization of a colocalization of IL-1R1 with SERT on serotonergic axons in this region supports this possibility. Our chronoamperometry studies were conducted in the dorsal CA3 which receives serotonergic input from both the DRN and MRN (57, 58). Although the MRN was not the focus of our current studies, we do detect expression of IL-1R1 by a significant subset of serotonergic neurons in this area (data not shown).

The redundancy in the effects that major proinflammatory cytokines have on cells has been observed in a multitude of body systems as part of a robust and reliable immune response (59, 60). The same appears to be in play in our studies with the effect that IL-1β and TNFα have on the excitability of DRN^5-HT^ neurons. We previously demonstrated a divergence in the effect produced by these cytokines on SERT, where both IL-1β and TNFα cause a p38α MAPK-dependent increase in SERT intrinsic activity, with TNFα acting through an additional p38α-independent manner to increase transporter surface trafficking (9), the basis for which has yet to be elucidated. Regardless, both cytokines exert an inhibitory effect on DRN^5-HT^ neuron excitability. Others have suggested that such redundancy may be important to ensure sufficient changes in target responses during acute inflammation, where levels of cytokines may not saturate their receptors (61). Alternatively, redundancy may be required to allow cytokines to access the same signaling cascade (e.g. p38α MAPK dependent pathways) if one cytokine precedes the other and wanes early as inflammation becomes more chronic. Future studies of how acute and chronic inflammation impact serotonergic signaling will be essential in order to develop therapeutics for individuals diagnosed with neuropsychiatric disorders who often display evidence of chronic low level inflammation. The common role of p38α MAPK in IL-1β and TNFα driven responses suggest that the kinase may be a more beneficial target for medication development in relation to CNS 5-HT linked disorders than either receptor alone.

To evaluate whether serotonergic IL-1R1 influences DRN^5-HT^ neuron activation *in vivo* as well as the forebrain targets of these cells, we pursued cFos studies as a reporter for neuronal activation. Indeed, serotonergic IL-1R1 was found to influence cFos labeling, with LPS administration decreasing the number of cFos+ serotonergic neurons of multiple DRN subregions. These findings are in line with our electrophysiological data in acute midbrain slices that demonstrate that the ability of IL1β (and TNFα) to reduce DRN^5-HT^ excitability. However, our results appear contrary to a prior study examining the effect of peripheral inflammation on 5-HT neuron activity as monitored using cFos (62). Hollis et al. performed similar acute LPS injections, although the dose was two times higher than ours, with findings of increased cFos labeling of 5-HT neurons within the dorsolateral DRN. Other differences between our two studies include mouse strain and age, with our mice being older adults. It has been shown that BALB/c mice, which were used in the Hollis study, have an elevated inflammatory response in comparison to C57BL/6, the background of the mice used in our study (63). Additional studies are needed to examine strain-related differences to immune stimuli in relation to regulation of 5-HT signaling *in vivo*.

When DRN subregions were examined for cFos responses to acute LPS, sex-specific differences emerged, suggesting that the forebrain target projections of DRN and/or MRN 5-HT neurons are also likely to display unique responses to inflammation. Indeed, our examination of one such projection target, the lateral habenula, showed a sex-specific decrease in cFos staining after LPS for the females and a nonsignificant increase in cFos staining in the males, with both effects normalized by the loss of serotonergic IL-1R1. The lateral habenula is a known forebrain target of DRN^5-HT^ neurons located in the dorsolateral DRN (50). Although activation within the lateral habenula is characteristic of negative reinforcement and depressive behavior, our acute inflammatory treatment may be showing an initiating and possibly transient state of lateral habenular neuronal inhibition (64, 65). Indeed, previous evidence shows that bath applied 5-HT activates the majority of lateral habenula neurons (66). Thus, an overall decrease in 5-HT signaling is expected to decrease the excitability of lateral habenular neurons. There is also a bidirectional communication between the DRN^5-HT^ neurons and the lateral habenula that should be taken into consideration when examining how this target region may be impacted by peripheral inflammation as well as inflammation associated neuropsychiatric disorders (50, 67). Glutamatergic neurons within the lateral habenula, specifically the medial portion of the lateral habenula project to the DRN (68). Electrical stimulation of the lateral habenula has been shown to suppress DRN^5-HT^ neurons, although this is likely an indirect effect with the lateral habenular glutamatergic neurons synapsing onto local DRN GABA neurons to elicit the effect (69, 70). Thus, an overall impact on 5-HT neuron activity is likely to have downstream effects that ultimately loop back to further influence DRN physiology. Approaches that allow for monitoring of both regions simultaneously in the context of inflammation are likely to be needed to understand the various consequences.

It appears that peripheral LPS elicits transcriptional changes in DRN^5-HT^ neurons not only in the form of cFos changes, but also in a multitude of other pathways. Using the GeoMx DSP spatial transcriptomics platform, we were able to pursue a spatial transcriptional analysis in a way that divided cell populations between high expressing IL-1R1 and low expressing IL-1R1 subregions of the DRN. In this report, we describe the effect of IL-1R1 on the more highly IL-1R1 expressing 5-HT neurons, with responses in the vDRN to be reported at a later time. Not only did we uncover a male-enriched transcriptional response to peripheral inflammation compared to females, we also found that the DEGs binned into sex-specific, nonoverlapping pathways. In the males, most of the DEGs were found to associate with RNA processing, translation, and protein modifications which suggests that male 5-HT neurons undergo broad alterations in fundamental cellular machinery, potentially reflecting a generalized stress-adaptive program. Inflammation-driven shifts in ribosomal and translational pathways have been found in a multitude of cell types, as the cells prepare to buffer against metabolic stress and promote proteostasis under inflammatory conditions (71, 72). On the other hand, females DEGs were found to be related to synaptic function, signaling and transport which suggests that female 5-HT neurons remodel neuronal communication and connectivity rather than core translational processes. Previous studies have shown that female brains often mount synapse-focused responses to immune challenges, leading to changes in plasticity and neurotransmission (73, 74). This suggests that male and female 5-HT neurons adapt to peripheral inflammation using different mechanisms, which may speak to the downstream behavioral differences noted in both inflammatory stimuli and depression between sexes (75, 76). The divergence may contribute to sex-specific vulnerabilities in neuropsychiatric disorders with inflammatory components, such as depression and anxiety, which are more prevalent in women and involve altered serotonergic signaling (77, 78).

Overall, this study expands our understanding of how inflammation, both peripheral and central, impacts the signaling capacity of serotonergic neurons via an IL-1R1 and p38α MAPK dependent pathway. Our work adds to the growing body of literature examining the interplay of 5-HT and neuroinflammation as a potential therapeutic target to alleviate neuroinflammatory symptoms in neuropsychiatric disorders, and recommends further evaluation of selective inhibitors of p38α MAPK such as MW150 (33) in the treatment of 5-HT associated brain disorders. Although our study did not include behavioral analyses, 5-HT is known to modulate a multitude of behaviors in rodent models and human. It is likely that the alterations in mood, appetite, and arousal that are seen in both infection and in those diagnosed with neurobehavioral disorders may be a result of the actions of inflammatory signals targeting of serotonergic neurons, where we demonstrate readily detectible expression of receptors for IL-1β and TNFα, two cytokines already shown to increase SERT activity. Future studies will examine how loss of serotonergic IL-1R1 impacts behavior and whether targeting inflammation is sufficient to restore altered serotonergic signaling that occurs in the context of neurobehavioral disorders.

